# TRACKing Tandem Repeats: a customizable pipeline for identification and cross-species comparisons

**DOI:** 10.1101/2024.09.27.615531

**Authors:** Carolina L. Adam, Joana Rocha, Peter Sudmant, Rori Rohlfs

**Affiliations:** Institute of Ecology and Evolution, University of Oregon, Eugene, OR 97403, USA; Department of Integrative Biology, University of California Berkeley, Berkeley, CA 94720, USA; School of Computer and Data Sciences, University of Oregon, Eugene, OR 97403, USA

**Keywords:** Tandem repeat variants, Comparative genomics, Evolution, Genomics, Population genetics

## Abstract

**Summary:** TRACK is a user-friendly command-line pipeline designed to consolidate the discovery and comparison of tandem repeats (TRs) across species. TRACK facilitates the cataloging and filtering of TRs based on reference genomes or T2T transcripts, and applies reciprocal LiftOver and sequence alignment methods to identify putative homologous TRs between species. For further streamlined analyses, TRACK can be used to genotype TRs and subsequently estimate and plot basic population genetic statistics. By integrating existing tools into one integrated workflow, TRACK enhances TR analysis accessibility and reproducibility, while offering flexibility for the user.

**Availability:** The TRACK toolkit with step-by-step tutorial is freely available at https://github.com/caroladam/track.

## 1. Background

Tandem repeats (TRs) are repetitive genomic sequences characterized by their abundance in genomes, high mutation rates, and presence of multiple alleles within a population (Gymrek, 2017), making them a major source of genetic variation (Kashi et al., 1997). By accumulating mutations faster than single nucleotide polymorphisms (SNPs) (Sun et al., 2012), TRs are an easy target for natural selection and act as central players in rapid evolution (Gemayel et al., 2010). Although ubiquitous in all eukaryotic genomes (Richard et al., 2008), TRs gained prominence in human and non-human primate studies due to their role as epigenetic and gene expression modulators and their association with many human diseases (Gymrek et al., 2016). TR expansions, for instance, are linked to the pathogenesis of multiple types of cancer (Erwin et al., 2023) and neurological-related conditions, such as Huntington’s disease (MacDonald et al., 1993), Friedreich’s ataxia (Campuzano et al., 1996), and psychiatric disorders (Song et al., 2018).

The cumulative evidence that TRs are associated with the evolution of complex traits (Press et al., 2014) and likely evolved under selective pressures (Liang et al., 2015) highlights the importance of understanding their variation across species through comparative analyses. In great apes, for example, there is evidence for the role of TRs in chromosomal rearrangements (Farré et al., 2011) and gene expression divergence (Bilgin Sonay et al., 2015). Thus, building comparative TR frameworks can provide critical insights into evolutionary processes.

The propensity of TRs for homoplasy, coupled with sequencing technology limitations that hindered accurate sequencing of long TRs, has historically restricted their widespread use in cross-species comparative studies (Hodel et al., 2016). The advent of long-read sequencing technologies provides the means to overcome these restrictions (Logsdon et al., 2020). The Telomere-to-Telomere (T2T) Consortium delivered the first gapless human genome assembly, CHM13, correcting inaccuracies in GRCh38 (Nurk et al., 2022), and has since expanded to include six non-human primate genomes (Yoo et al., 2024). These methods are now being applied to generate T2T genomes for additional taxa across eukaryotes (Kalbfleisch et al., 2024; Zhang et al., 2023). These high-resolution data and the availability of new analysis tools tailored for TRs (Dolzhenko et al., 2024; Mousavi et al., 2020) open avenues for comparative studies with unparalleled resolution, enabling the discovery of previously undetected TRs, which may help generate new evolutionary hypotheses.

To simplify the discovery and analysis of shared TR loci across species, we developed the **T**andem **R**epeat **A**nalysis and **C**omparison **K**it (**TRACK**). TRACK leverages established analytical tools to efficiently identify TR loci from chromosome-level genome assemblies, producing comprehensive catalogs of putative homologous TRs between species. Beyond cataloging, TRACK streamlines the process of genotyping these TRs in population-level datasets, offering a suite of options to calculate and visualize key population genetics estimates (**Figure 1**). This all-in-one toolkit provides a unified approach to uncovering evolutionary insights from TR data.

**Figure 1.**
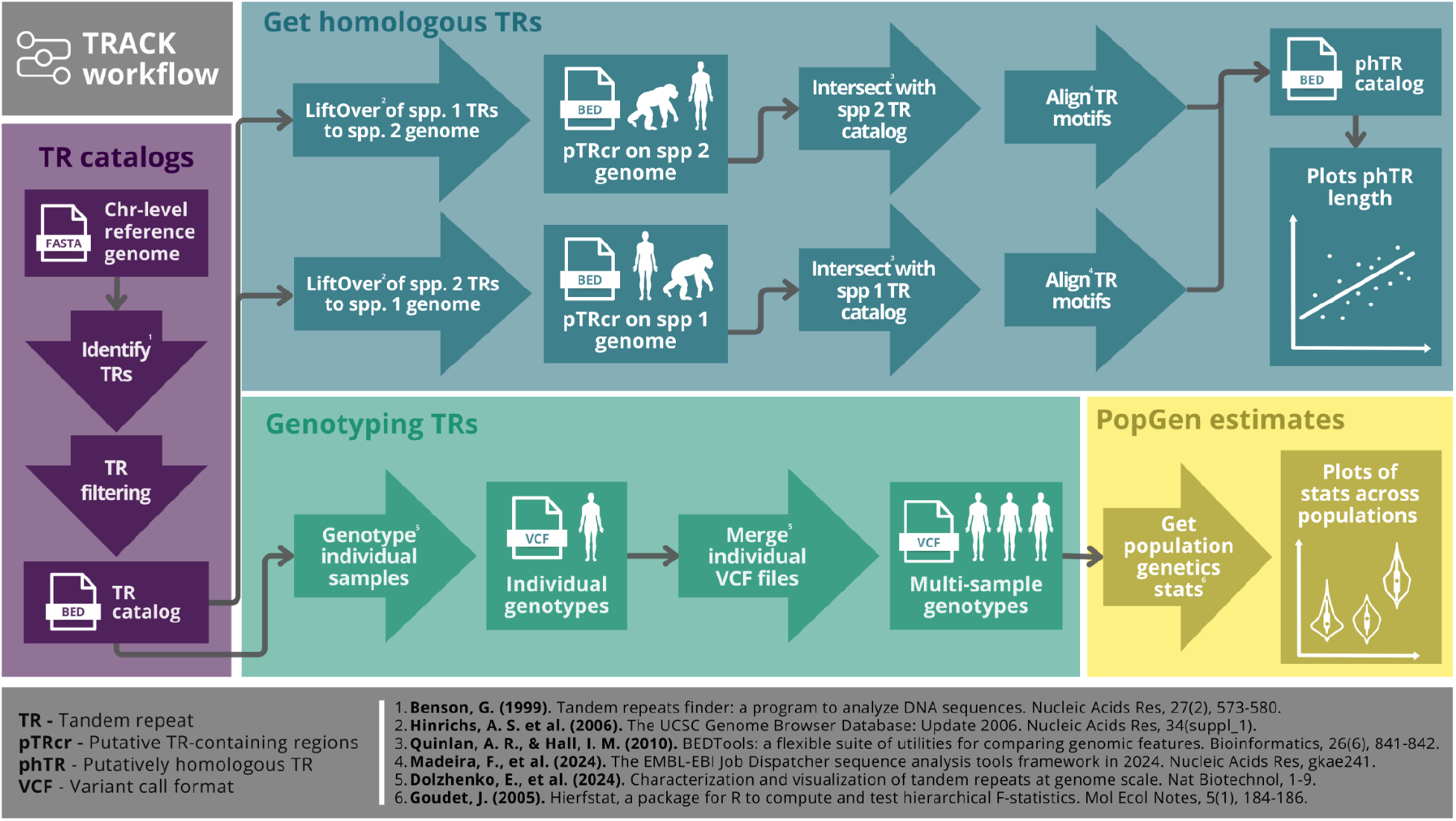
A simplified workflow of the Tandem Repeat Analysis and Comparison Kit (TRACK) features.

## 2. Pipeline description

### 2.1 Generating TR catalogs

TRACK uses Tandem Repeat Finder (TRF) version 4.09 (Benson, 1999) to generate the TR reference catalogs from chromosome-level reference genomes. TRACK default parameters are in Table 1, which results in catalogs with TR minimum length >12bp. To eliminate redundancy from multiple computations at the same genomic index position or variation in score values, overlapping TRs are initially merged. Subsequently, TRs exceeding 10 Kbp in total length or having a copy number <2.5 are filtered out.

**Table 1.**
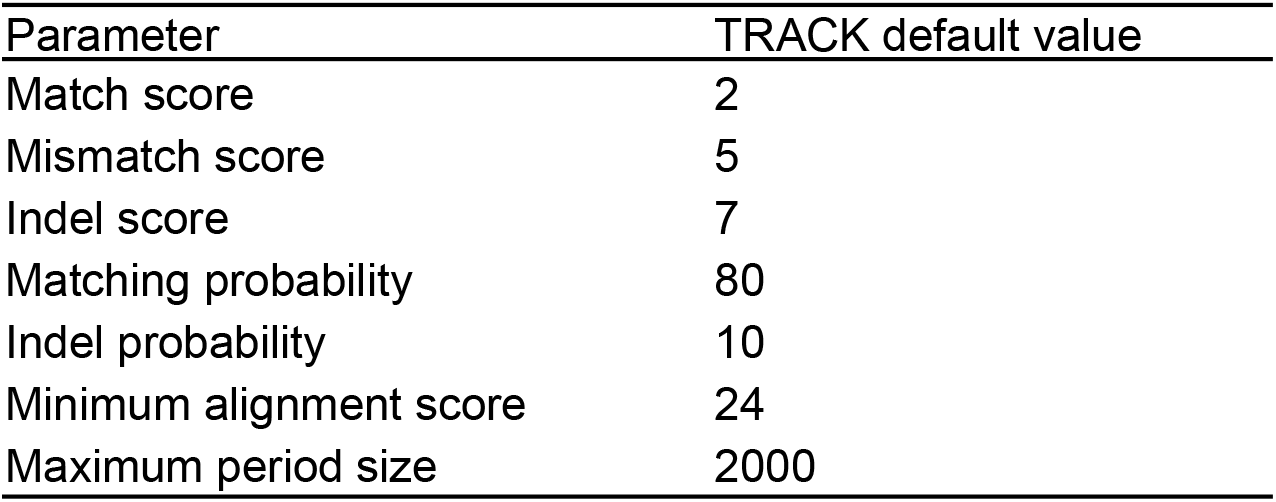
TRACK default parameters for Tandem Repeat Finder (TRF) run.

| Parameter | TRACK default value |
| --- | --- |
| Match score | 2 |
| Mismatch score | 5 |
| Indel score | 7 |
| Matching probability | 80 |
| Indel probability | 10 |
| Minimum alignment score | 24 |
| Maximum period size | 2000 |

### 2.2 Identification of putatively homologous TRs

Putative TR homology is initially assessed in a pairwise manner. TRACK begins by conducting a LiftOver (Hinrichs et al., 2006) analysis using a TR reference catalog from a target genome (tTRc) and a chain file that describes the conversion between genome positions from a target genome assembly to a query genome assembly. Chain files can be obtained directly from the UCSC Genome Browser or be custom-made by the user by performing whole-genome alignment of target and query genomes and subsequent conversion of the alignment file to chain file. The liftover step produces a file of putative TR-containing regions (pTRcr) in the query genome. To reduce bias in homology detection, TRACK performs the LiftOver analysis bidirectionally, yielding two lifted *bed* files for each pairwise comparison. The resulting pTRcr file is then compared to the original TR reference catalog of the query genome, retaining only regions that meet a user-defined reciprocal overlap threshold. This means that both the lifted TR region and the corresponding TR in the reference catalog must overlap by the specified percentage to be considered putatively homologous.

To verify the motif similarity of these TRs based on sequence composition, TRACK conducts pairwise global alignments between the TR motifs from each species using the Needleman-Wunsch algorithm (Madeira et al., 2024). Then, TRACK only keeps regions that meet a user-specified threshold for TR motif sequence similarity. The two resulting files from each pairwise comparison are then intersected to produce a catalog of putatively homologous TRs (phTRs), one per row, in .bed format. TRACK also provides a visualization feature to create a scatterplot comparing the total length of shared TRs between two individuals of different species (**Figure 2**).

**Figure 2.**
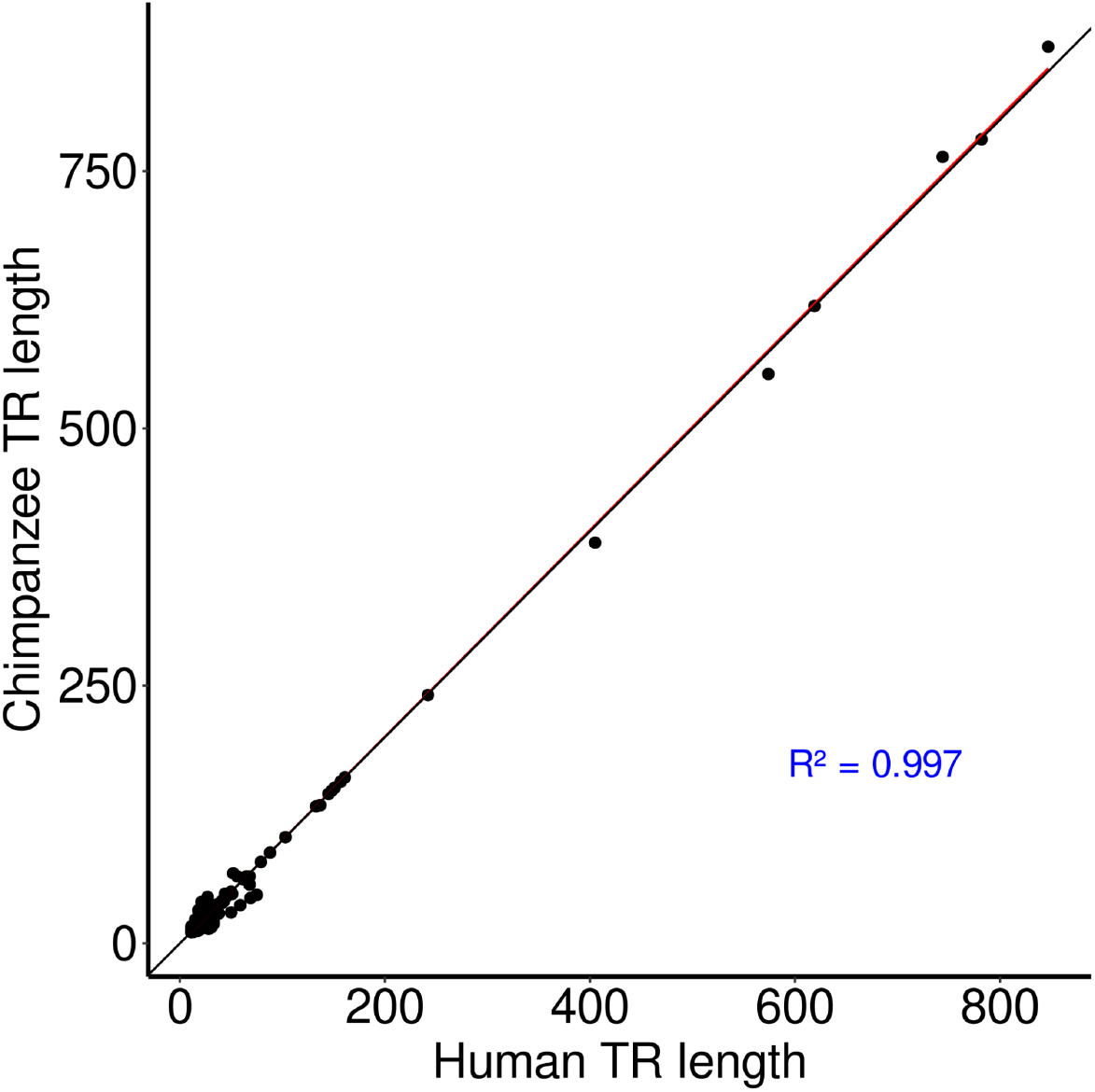
Scatterplot of the total length of a subset of 1,000 TRs shared between human and chimpanzee T2T reference genomes.

### 2.3 Genotyping TRs and basic population genetic statistics

One of the many uses of TR catalogs is genotyping variants in within-species population datasets. TRACK provides an integrated module for preparing and structuring long-read data to genotype TR variants using the Tandem Repeat Genotyping Tool (TRGT) (Dolzhenko et al., 2024). The output is a VCF file containing genotyped TR variants across multiple samples. Additionally, TRACK provides scripts to estimate basic population genetic statistics, such as observed heterozygosity and genetic diversity, and generates violin plots to visualize the results (**Figure 3**).

**Figure 3.**
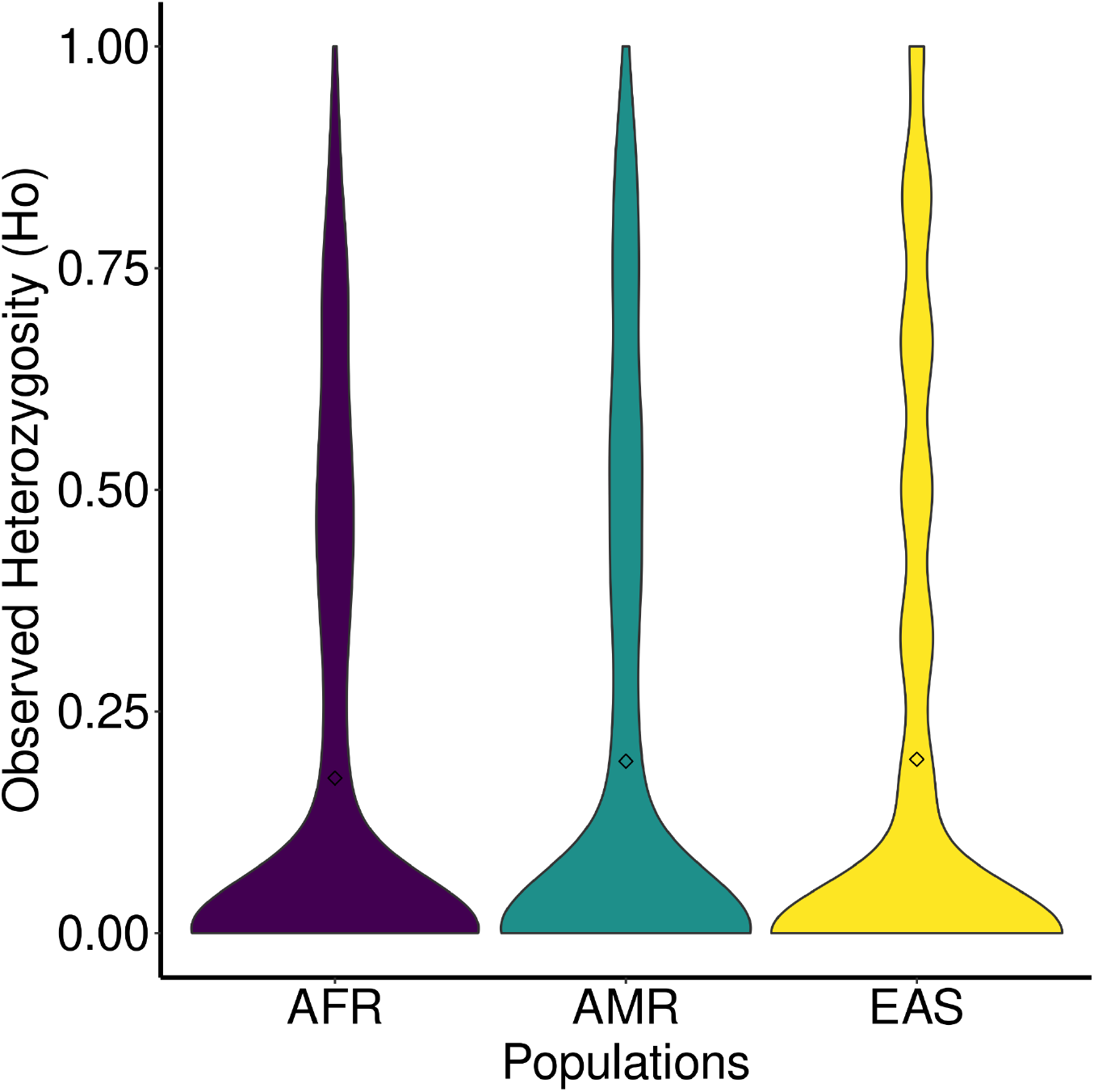
Observed heterozygosity estimates from a random subset of 1,000 TR loci from TRACK’s human T2T TR catalog genotyped in individuals of the Human Pangenome Reference Consortium (HPRC) (Wang et al., 2022). AFR - Africa; AMR - Americas; EAS - Asia.

## 3. Conclusion

The significance of TRACK lies in simplifying and streamlining some of the often complex and time-consuming steps of comparative TR analysis. By integrating multiple features from different established tools—from catalog generation and cross-species comparison to population-level genotyping and diversity estimates—TRACK improves the accessibility of TR analyses while increasing reproducibility across analyses. Additionally, TRACK is highly flexible, allowing the user to start from different points of the pipeline. This is particularly valuable as the volume of genomic data grows, making tools like TRACK essential for facilitating the discovery of biologically meaningful insights about TRs from long-read sequence data.

## REFERENCES

Benson, G. (1999). Tandem repeats finder: A program to analyze DNA sequences. Nucleic Acids Research, 27(2), 573–580. 10.1093/nar/27.2.573

Bilgin Sonay, T., Carvalho, T., Robinson, M. D., Greminger, M. P., Krützen, M., Comas, D., Highnam, G., Mittelman, D., Sharp, A., Marques-Bonet, T., & Wagner, A. (2015). Tandem repeat variation in human and great ape populations and its impact on gene expression divergence. Genome Research, 25(11), 1591–1599. 10.1101/gr.190868.115

Campuzano, V., Montermini, L., Moltò, M. D., Pianese, L., Cossée, M., Cavalcanti, F., Monros, E., Rodius, F., Duclos, F., Monticelli, A., Zara, F., Cañizares, J., Koutnikova, H., Bidichandani, S. I., Gellera, C., Brice, A., Trouillas, P., De Michele, G., Filla, A., … Pandolfo, M. (1996). Friedreich’s Ataxia: Autosomal Recessive Disease Caused by an Intronic GAA Triplet Repeat Expansion. Science, 271(5254), 1423–1427. 10.1126/science.271.5254.1423

Dolzhenko, E., English, A., Dashnow, H., De Sena Brandine, G., Mokveld, T., Rowell, W. J., Karniski, C., Kronenberg, Z., Danzi, M. C., Cheung, W. A., Bi, C., Farrow, E., Wenger, A., Chua, K. P., Martínez-Cerdeño, V., Bartley, T. D., Jin, P., Nelson, D. L., Zuchner, S., … Eberle, M. A. (2024). Characterization and visualization of tandem repeats at genome scale. Nature Biotechnology, 1–9. 10.1038/s41587-023-02057-3

Erwin, G. S., Gürsoy, G., Al-Abri, R., Suriyaprakash, A., Dolzhenko, E., Zhu, K., Hoerner, C. R., White, S. M., Ramirez, L., Vadlakonda, A., Vadlakonda, A., Von Kraut, K., Park, J., Brannon, C. M., Sumano, D. A., Kirtikar, R. A., Erwin, A. A., Metzner, T. J., Yuen, R. K. C., … Snyder, M. P. (2023). Recurrent repeat expansions in human cancer genomes. Nature, 613(7942), 96–102. 10.1038/s41586-022-05515-1

Farré, M., Bosch, M., López-Giráldez, F., Ponsà, M., & Ruiz-Herrera, A. (2011). Assessing the Role of Tandem Repeats in Shaping the Genomic Architecture of Great Apes. PLoS ONE, 6(11), e27239. 10.1371/journal.pone.0027239

Gemayel, R., Vinces, M. D., Legendre, M., & Verstrepen, K. J. (2010). Variable Tandem Repeats Accelerate Evolution of Coding and Regulatory Sequences. Annual Review of Genetics, 44(1), 445–477. 10.1146/annurev-genet-072610-155046

Gymrek, M. (2017). A genomic view of short tandem repeats. Current Opinion in Genetics & Development, 44, 9–16. 10.1016/j.gde.2017.01.012

Gymrek, M., Willems, T., Guilmatre, A., Zeng, H., Markus, B., Georgiev, S., Daly, M. J., Price, A. L., Pritchard, J. K., Sharp, A. J., & Erlich, Y. (2016). Abundant contribution of short tandem repeats to gene expression variation in humans. Nature Genetics, 48(1), 22–29. 10.1038/ng.3461

Hinrichs, A. S., Karolchik, D., Baertsch, R., Barber, G. P., Bejerano, G., Clawson, H., Diekhans, M., Furey, T. S., Harte, R. A., Hsu, F., Hillman-Jackson, J., Kuhn, R. M., Pedersen, J. S., Pohl, A., Raney, B. J., Rosenbloom, K. R., Siepel, A., Smith, K. E., Sugnet, C. W., … Kent, W. J. (2006). The UCSC Genome Browser Database: Update 2006. Nucleic Acids Research, 34(suppl_1), D590–D598. 10.1093/nar/gkj144

Hodel, R. G. J., Segovia-Salcedo, M. C., Landis, J. B., Crowl, A. A., Sun, M., Liu, X., Gitzendanner, M. A., Douglas, N. A., Germain-Aubrey, C. C., Chen, S., Soltis, D. E., & Soltis, P. S. (2016). The Report of My Death was an Exaggeration: A Review for Researchers Using Microsatellites in the 21st Century. Applications in Plant Sciences, 4(6), 1600025. 10.3732/apps.1600025

Kalbfleisch, T. S., McKay, S. D., Murdoch, B. M., Adelson, D. L., Almansa-Villa, D., Becker, G., Beckett, L. M., Benítez-Galeano, M. J., Biase, F., Casey, T., Chuong, E., Clark, E., Clarke, S., Cockett, N., Couldrey, C., Davis, B. W., Elsik, C. G., Faraut, T., Gao, Y., … Rosen, B. D. (2024). The Ruminant Telomere-to-Telomere (RT2T) Consortium. Nature Genetics, 56(8), 1566–1573. 10.1038/s41588-024-01835-2

Kashi, Y., King, D., & Soller, M. (1997). Simple sequence repeats as a source of quantitative genetic variation. Trends in Genetics, 13(2), 74–78. 10.1016/S0168-9525(97)01008-1

Liang, K.-C., Tseng, J. T., Tsai, S.-J., & Sun, H. S. (2015). Characterization and distribution of repetitive elements in association with genes in the human genome. Computational Biology and Chemistry, 57, 29–38. 10.1016/j.compbiolchem.2015.02.007

Logsdon, G. A., Vollger, M. R., & Eichler, E. E. (2020). Long-read human genome sequencing and its applications. Nature Reviews Genetics, 21(10), 597–614. 10.1038/s41576-020-0236-x

MacDonald, M. E., Ambrose, C. M., Duyao, M. P., Myers, R. H., Lin, C., Srinidhi, L., Barnes, G., Taylor, S. A., James, M., Groot, N., MacFarlane, H., Jenkins, B., Anderson, M. A., Wexler, N. S., Gusella, J. F., Bates, G. P., Baxendale, S., Hummerich, H., Kirby, S., … Harper, P. S. (1993). A novel gene containing a trinucleotide repeat that is expanded and unstable on Huntington’s disease chromosomes. Cell, 72(6), 971–983. 10.1016/0092-8674(93)90585-E

Madeira, F., Madhusoodanan, N., Lee, J., Eusebi, A., Niewielska, A., Tivey, A. R. N., Lopez, R., & Butcher, S. (2024). The EMBL-EBI Job Dispatcher sequence analysis tools framework in 2024. Nucleic Acids Research, 52(W1), W521–W525. 10.1093/nar/gkae241

Mousavi, N., Margoliash, J., Pusarla, N., Saini, S., Yanicky, R., & Gymrek, M. (2020). TRTools: A toolkit for genome-wide analysis of tandem repeats. Bioinformatics, 37(5), 731–733. 10.1093/bioinformatics/btaa736

Nurk, S., Koren, S., Rhie, A., Rautiainen, M., Bzikadze, A. V., Mikheenko, A., Vollger, M. R., Altemose, N., Uralsky, L., Gershman, A., Aganezov, S., Hoyt, S. J., Diekhans, M., Logsdon, G. A., Alonge, M., Antonarakis, S. E., Borchers, M., Bouffard, G. G., Brooks, S. Y., … Phillippy, A. M. (2022). The complete sequence of a human genome.

Press, M. O., Carlson, K. D., & Queitsch, C. (2014). The overdue promise of short tandem repeat variation for heritability. Trends in Genetics, 30(11), 504–512. 10.1016/j.tig.2014.07.008

Richard, G.-F., Kerrest, A., & Dujon, B. (2008). Comparative Genomics and Molecular Dynamics of DNA Repeats in Eukaryotes. Microbiology and Molecular Biology Reviews, 72(4), 686–727. 10.1128/MMBR.00011-08

Song, J. H. T., Lowe, C. B., & Kingsley, D. M. (2018). Characterization of a Human-Specific Tandem Repeat Associated with Bipolar Disorder and Schizophrenia. American Journal of Human Genetics, 103(3), 421–430. 10.1016/j.ajhg.2018.07.011

Sun, J. X., Helgason, A., Masson, G., Ebenesersdóttir, S. S., Li, H., Mallick, S., Gnerre, S., Patterson, N., Kong, A., Reich, D., & Stefansson, K. (2012). A direct characterization of human mutation based on microsatellites. Nature Genetics, 44(10), 1161–1165. 10.1038/ng.2398

Wang, T., Antonacci-Fulton, L., Howe, K., Lawson, H. A., Lucas, J. K., Phillippy, A. M., Popejoy, A. B., Asri, M., Carson, C., Chaisson, M. J. P., Chang, X., Cook-Deegan, R., Felsenfeld, A. L., Fulton, R. S., Garrison, E. P., Garrison, N. A., Graves-Lindsay, T. A., Ji, H., Kenny, E. E., … Haussler, D. (2022). The Human Pangenome Project: A global resource to map genomic diversity. Nature, 604(7906), 437–446. 10.1038/s41586-022-04601-8

Yoo, D., Rhie, A., Hebbar, P., Antonacci, F., Logsdon, G. A., Solar, S. J., Antipov, D., Pickett, B. D., Safonova, Y., Montinaro, F., Luo, Y., Malukiewicz, J., Storer, J. M., Lin, J., Sequeira, A. N., Mangan, R. J., Hickey, G., Anez, G. M., Balachandran, P., … Eichler, E. E. (2024). Complete sequencing of ape genomes. 10.1101/2024.07.31.605654

Zhang, C., Xie, L., Yu, H., Wang, J., Chen, Q., & Wang, H. (2023). The T2T genome assembly of soybean cultivar ZH13 and its epigenetic landscapes. Molecular Plant, 16(11), 1715–1718. 10.1016/j.molp.2023.10.003

